# *Prepronociceptin*-expressing neurons in the extended amygdala signal darting away from an aversive odor

**DOI:** 10.1101/2022.02.21.481217

**Authors:** Randall L. Ung, Maria M. Ortiz-Juza, Vincent R. Curtis, Rizk A. Alghorazi, Geronimo Velazquez-Hernandez, Ayden Ring, Ruben A. Garcia-Reyes, Garret D. Stuber, Pengcheng Zhou, Hiroyuki K. Kato, Nicolas C. Pégard, Jose Rodriguez-Romaguera

## Abstract

Dysregulation in the neural circuitry that encodes physiological arousal responses is thought to contribute to the manifestation of the maladaptive behaviors observed in neuropsychiatric disorders. We previously found that *prepronociceptin*-expressing neurons in the bed nucleus of the stria terminalis (*Pnoc*^BNST^ neurons) modulate rapid changes in physiological arousal upon presentation of motivationally salient stimuli (Rodriguez-Romaguera et al., 2020). However, whether *Pnoc*^BNST^ neurons are necessary to regulate behavioral actions to motivationally salient stimuli is still unknown. Here, we investigated the role of *Pnoc*^BNST^ neurons in encoding behavioral responses to motivationally salient stimuli using *in vivo* calcium imaging and optogenetic approaches in freely behaving mice. We find that the bulk activity of *Pnoc*^BNST^ neurons increases when mice are near an aversive odor in comparison to a rewarding odor. However, optogenetic inhibition of *Pnoc*^BNST^ neurons does not affect the amount of time mice spend near an aversive odor. Further analysis revealed that a subgroup of *Pnoc*^BNST^ neurons that correlate with proximity to the aversive odor also correlate to darting away from the same aversive odor. Since these two behaviors are opposite to each other and since we previously found *Pnoc*^BNST^ neurons correlate with arousal responses, we believe these results may be due in part to the encoding of arousal responses that occur when mice approach and dart away from aversive stimuli.

## INTRODUCTION

Atypical physiological arousal responses are a core component of many neuropsychiatric disorders that are also characterized by maladaptive behavior. For instance, patients suffering from anxiety disorders experience heightened arousal responses to stimuli and display deficits in motivated behavior (Wilhelm and Roth, 2001; Maclean and Datta, 2007; Craske et al., 2009; Lang and McTeague, 2009; Schmidt et al., 2017; Urbano et al., 2017; Patriquin et al., 2019). This comorbidity suggests that common circuitry contributes to both these symptoms. Despite this association, many studies do not investigate the relationship between the circuitry that governs internal arousal responses with circuits that regulate motivated behavioral states. Therefore, new experiments that evaluate how the neuronal ensembles that regulate arousal responses contribute to ongoing motivated behavior can elucidate the circuit-based mechanisms of how changes within internal processes impact external behaviors.

The bed nucleus of the stria terminalis (BNST) is a genetically diverse hub of the extended amygdala (Giardino and Pomrenze, 2021; Ortiz-Juza et al., 2021), and is strongly implicated in human psychopathologies defined by alterations in arousal and motivational states (Straube et al., 2007; Walker et al., 2009; Yassa et al., 2012). Recently, we identified that the activity of a subpopulation of *prepronociceptin*-expressing neurons within the BNST (*Pnoc*^BNST^ neurons) directly correlates with rapid changes in physiological arousal (observed by increases in pupillary size) when mice are presented with motivationally salient predator and food odors (Rodriguez-Romaguera et al., 2020). In addition, optogenetic activation of *Pnoc*^BNST^ neurons induced rapid increases in physiological arousal, indicating that *Pnoc*^BNST^ neurons are necessary for encoding these responses to motivationally salient stimuli. However, whether the response dynamics of *Pnoc*^BNST^ neurons are also necessary for encoding the behavioral actions that occur in response to motivationally salient stimuli remains unknown.

In the present study, we performed a series of *in vivo* calcium imaging and optogenetic studies in freely behaving mice to assess the necessity of *Pnoc*^BNST^ neurons in encoding motivated behavioral actions to aversive and rewarding odors. We found that the bulk activity of *Pnoc*^BNST^ neurons significantly increases when mice are near an aversive odor, as compared to a rewarding odor. Bulk inhibition of *Pnoc*^BNST^ neurons does not influence the amount of time mice spend near an aversive odor. However, when we define *Pnoc*^BNST^ neuronal activity by their individual activity dynamics, we find that an ensemble of neurons that was significantly excited when mice went near the aversive odor for the first time, subsequently correlated with darting away from the same aversive odor. Therefore, our findings reveal that *Pnoc*^BNST^ neurons signal the action to dart away from a motivationally salient stimulus with a negative valence.

## RESULTS

### Distinct motivationally salient odors induce approach and avoidance behavior

The BNST coordinates multiple motivational states that are essential for guiding actions to seek reward and/or avoid aversive stimuli. We previously observed, in head-fixed and restrained mice, that the activity of *Pnoc*^BNST^ neurons showed a high degree of correlation with changes in internal physiologic responses (pupillary dynamics) upon the presentation of both aversive and rewarding odors (Rodriguez-Romaguera et al., 2020). However, whether these neural dynamics, which are associated with arousal responses, signal motivated actions towards or away from salient stimuli has not been studied. Here, we sought to understand the encoding properties of *Pnoc*^BNST^ neurons on the behavioral actions that mice exhibit when exposed to arousal-inducing odors of opposing valence: appetitive odor - peanut oil, and aversive odor - trimethylthiazoline (TMT). To assess approach and avoidance behavior to these salient stimuli, we exposed mice to an odor preference assay within their home cage to maintain a familiar environment and avoid behavior associated with a novel context. We determined a baseline side preference by exposing mice to a control odor swab that contained water for 5 minutes and measured the time spent in far, transition, and near zones relative to the location of the odor swab. This was followed by 5 minutes of exposure to either a peanut swab (placed in the non-preferred zone) or a TMT swab (placed in the preferred zone) **(Figure 1A and 1B)**. We found that mice exhibit a significant preference for the peanut swab, as indicated by increased time spent in the near zone compared to the control swab **(Figure 1C)**. In addition, we found that mice exhibit significant aversion to the TMT swab, as indicated by decreased time spent in the near zone compared to the control swab **(Figure 1D)**. These results demonstrate that peanut and TMT odors, respectively, induce approach and avoidance behaviors in an odor preference assay.

**Figure 1.**
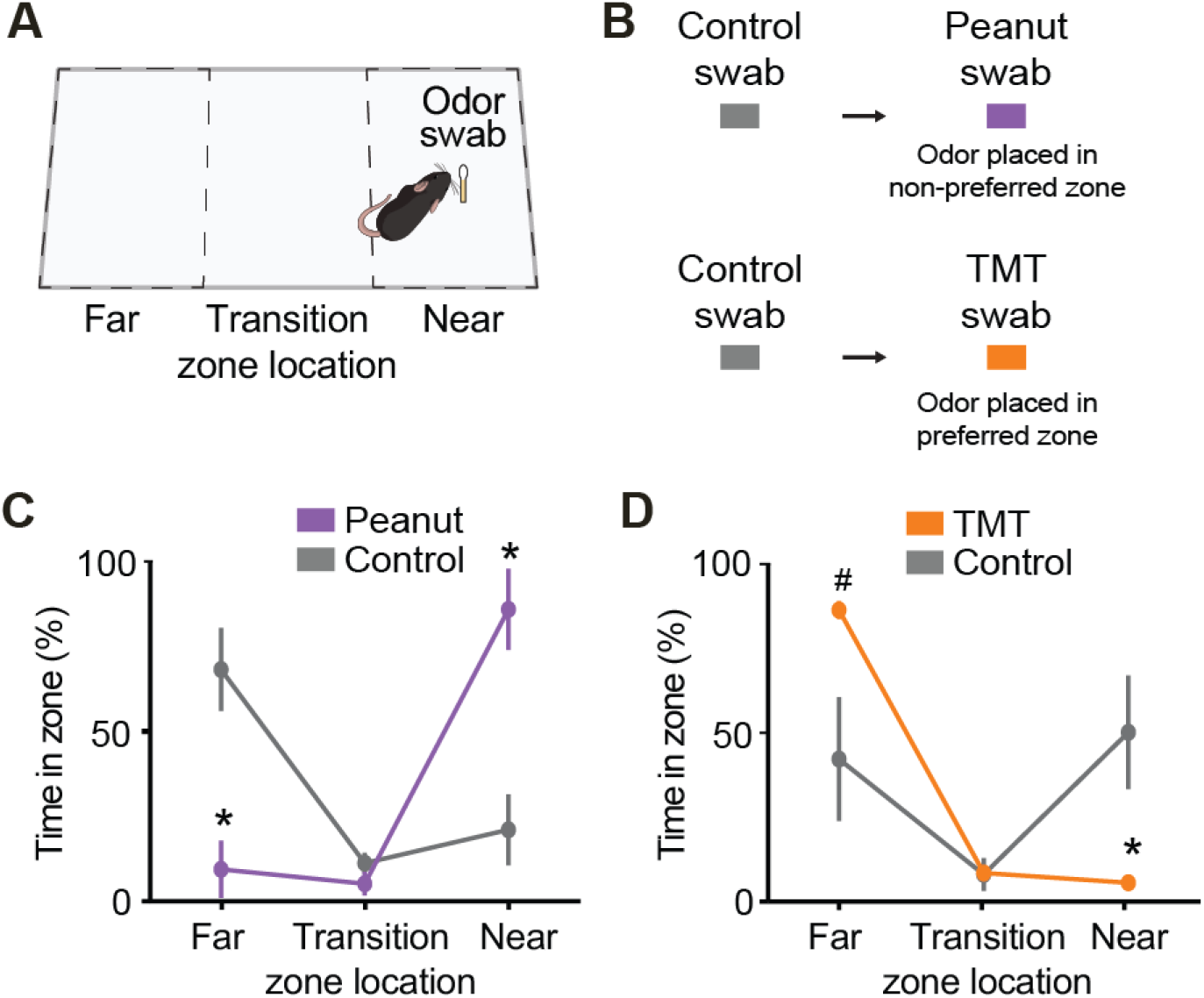
Distinct motivationally salient odors induce approach or avoidance behaviors. **(A)** Schematic of a freely moving mouse in its home cage. We assess approach and avoidance behaviors to water (2.5 μl), a low dose (2.5 μl) of 2,3,5-trimethylthiazoline (TMT), or peanut oil (2.5 μl) via a cotton swab placed on a platform on one side of the home cage. **(B)** Schematic showing order of control and odor swab presentation. **(C)** Group average of time spent across zones relative to the location of odor source in mice exposed to control or peanut oil swabs (n= 5 *Pnoc*-IRES-Cre mice). **(D)** Group average of time spent across zones relative to the location of odor source in mice exposed to control or TMT oil swabs (n=6 *Pnoc*-IRES-Cre mice). Data are shown as mean ± SEM. *\*p<0.05, ^#^p<0.1*.

### Aversive, but not rewarding odors, increase the bulk activity of *Pnoc*^BNST^ neurons

We previously observed that the activity of *Pnoc*^BNST^ neurons showed a high degree of correlation with physiological arousal responses upon the presentation of both aversive and rewarding odors in head-fixed mice (Rodriguez-Romaguera et al., 2020). To test if the activity of *Pnoc*^BNST^ neurons also correlates with avoidance and approach behaviors to these same odors, we performed *in vivo* calcium imaging in freely behaving mice. We used a head-mounted miniature fluorescence microscope (Ghosh et al., 2011; Resendez et al., 2016; Aharoni and Hoogland, 2019) to visualize the neural dynamics of *Pnoc*^BNST^ neurons that were transduced to express GCaMP6s (see Surgical Procedure and Histology methods section) **(Figure 2A-C)**. After habituation to the microscope, mice were exposed to water, peanut oil, or TMT on a cotton swab that was positioned on one side of their home cage using a square block holder that maintained a 25 cm maximum separation between the mouse and odor swab. Here, we analyzed the average calcium activity of *Pnoc*^BNST^ neurons while mice explored the far, transition, or near zones relative to the odor source **(Figure 2D-E)**. Group analysis of the calcium activity of *Pnoc*^BNST^ neurons showed no difference when mice were presented with peanut oil in their home cage compared to the water swab and this remained consistent across all zone locations **(Figure 2F)**. However, bulk analysis of the calcium activity of *Pnoc*^BNST^ neurons when mice were presented with TMT in their home cage showed a significant increase in calcium activity compared to the water swab with the greatest increase of activity observed when mice were located near the TMT odor source **(Figure 2G)**. We also found that when we averaged the calcium activity of *Pnoc*^BNST^ neurons across all zones and compared it between the two odor sources, TMT induced a greater change in calcium activity compared to peanut oil **(Figure 2H)**. These results suggest that *Pnoc*^BNST^ neurons may play a role in encoding the behavioral actions that occur in the presence of aversive odors.

**Figure 2.**
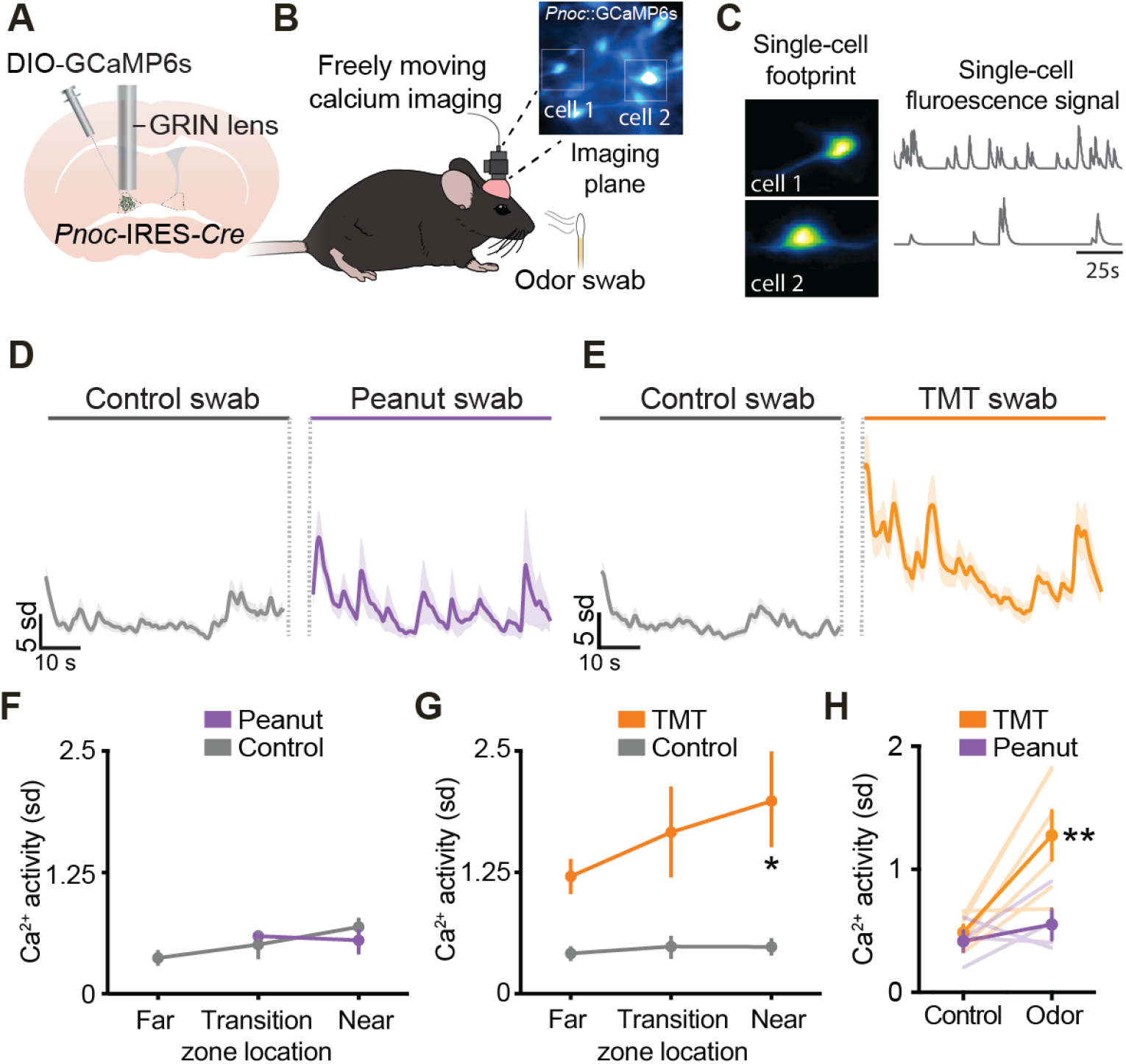
TMT, but not Peanut oil, increases bulk activity of *Pnoc*^BNST^ neurons. **(A)** Schematic of implantation of a GRIN lens above adBNST of *Pnoc*-IRES-*Cre* mice injected with AAVdj-EF1α-DIO-GCaMP6s. **(B)** *Left*: Schematic of a freely moving mouse with a head-mountable miniature microscope. *Right*: Representative image of *Pnoc*^BNST^ neurons through a GRIN lens. **(C)** Extracted calcium traces from two representative *Pnoc*^BNST^ neurons using CNMF. **(D)** Bulk response dynamics of *Pnoc*^BNST^ neurons to control swab and peanut swab. **(E)** Bulk response dynamics of *Pnoc*^BNST^ neurons to control swab and to TMT swab. **(F)** Comparison of the bulk response dynamics of *Pnoc*^BNST^ neurons of control swab and peanut swab across zones (n= 96 *Pnoc*^+^ neurons from 4 mice). **(G)** Comparison of the bulk response dynamics of *Pnoc*^BNST^ neurons of control swab and TMT swab across zones (n=163 *Pnoc*^+^ neurons from 6 mice). **(H)** Average bulk response dynamics of *Pnoc*^BNST^ neurons to peanut swab and TMT swab. Data are shown as mean ± SEM. *\*p<0.05, **p<0.01*.

### Bulk inhibition of *Pnoc*^BNST^ neurons does not prevent avoidance of TMT odor

Since we observed a significant increase in calcium activity of *Pnoc*^BNST^ neurons when mice were exposed to aversive odor TMT, we next tested whether photoinhibition of *Pnoc*^BNST^ neurons would change the amount of time mice spend near TMT odor. First, we evaluated whether viral tools can reliably inhibit *Pnoc*^BNST^ neuronal activity using a head-mountable miniature microscope capable of performing both *in vivo* calcium imaging and optogenetics. We unilaterally injected *Pnoc*-IRES-*Cre* mice with a 1:1 (v/v) viral solution that contained both AAVdj-EF1α-DIO-GCaMP6s and AAV5-EF1α-DIO-eNpHR3.0-mCherry into adBNST **(Figure 3A)**. We found that the photoinhibition of *Pnoc*^BNST^ neurons during a 3-min epoch significantly reduced calcium transients in freely moving animals and that the reduction in activity was solely observed during periods in which the optogenetic stimulation laser was on **(Figure 3B)**. We next tested if we could increase the time amount of time spent near TMT odor by photoinhibition of *Pnoc*^BNST^ neurons during 3-min laser on vs off epochs. We injected AAVs carrying EF1α-DIO-eNpHR3.0-eYFP into adBNST of either *Pnoc*-IRES-*Cre* mice or their wild-type littermates and implanted optic fibers above adBNST **(Figure 3C)**. Constant 532 nm green laser light was delivered for 3-min epochs while mice were exposed to TMT. We found that photoinhibition of *Pnoc*^BNST^ neurons did not change the amount of time mice spent near TMT odor **(Figure 3D)**. These results show that *Pnoc*^BNST^ neurons are not necessary for avoidance of TMT odor.

**Figure 3.**
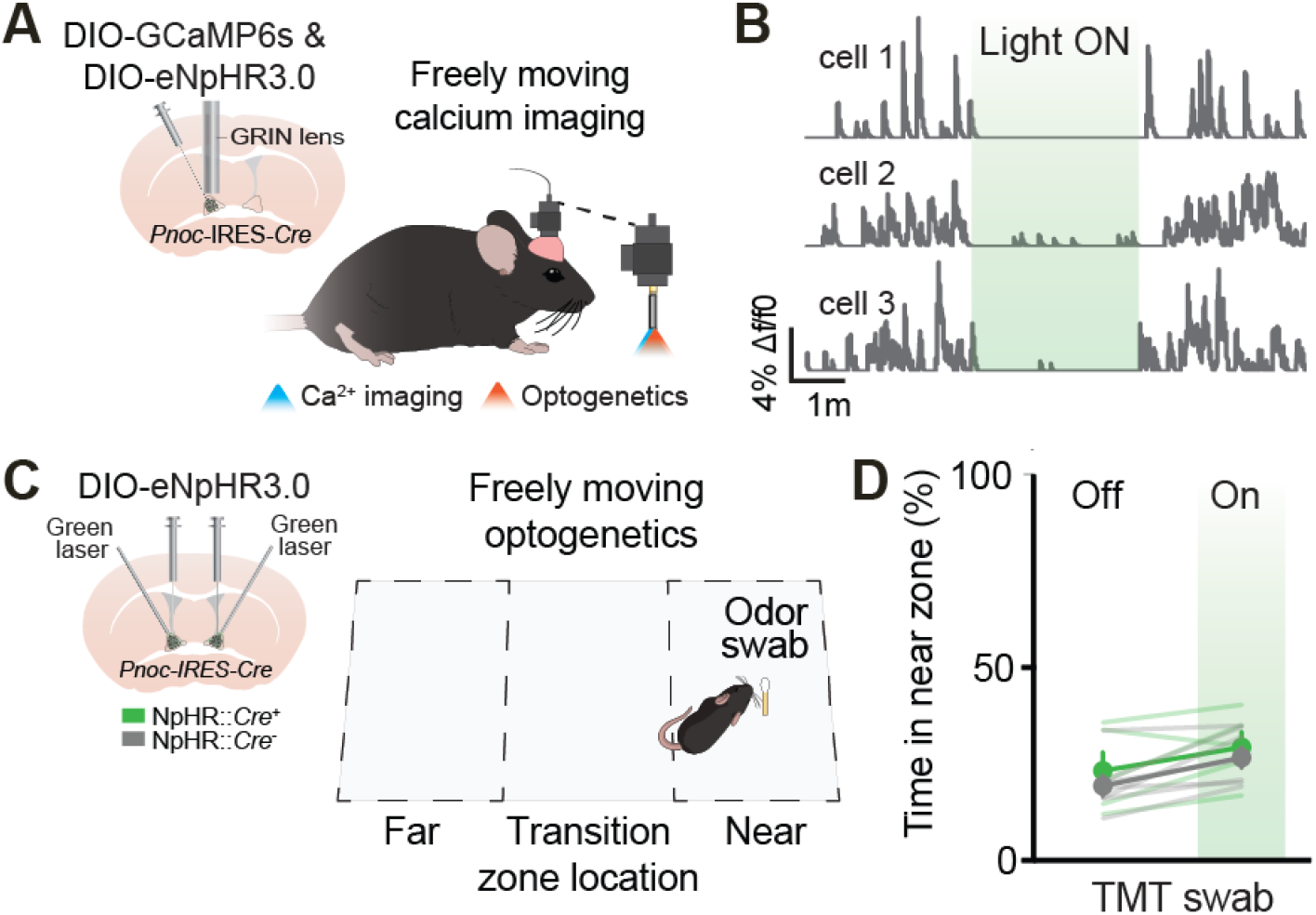
Bulk inhibition of *Pnoc*^BNST^ neurons does not prevent avoidance of TMT odor. **(A)** *Left*: Schematic of implantation of a GRIN lens above adBNST of *Pnoc*-IRES-*Cre* mice injected with AAVdj-EF1α-DIO-GCaMP6s & DIO-eNpHR3.0. *Right*: Schematic of a freely moving mouse with a head-mountable miniature microscope that allows for simultaneous calcium imaging and optogenetics. **(B)** Calcium traces from three representative *Pnoc*^BNST^ neurons before, during, and after optogenetic inhibition. **(C)** *Left*: Schematic of implantation of optical fibers implanted above adBNST of *Pnoc*-IRES-*Cre* mice injected with DIO-eNpHR3.0. *Right*: Schematic of a freely moving mouse in its home cage to assess approach and avoidance behaviors to TMT swab while inhibiting *Pnoc*^BNST^ neurons. **(D)** Grouped average of time spent near TMT during periods when the laser was off vs. on (n=7 NpHR-Cre^-^ mice, n=5 NpHR-Cre^+^ mice). Data is shown as mean ± SEM.

### Identifying neurons based on individual response dynamics to TMT and Peanut

Since bulk inhibition of *Pnoc*^BNST^ neurons did not alter avoidance behavior to TMT odor as we had predicted, we next performed analysis on the individual response dynamics of *Pnoc*^BNST^ neurons when mice were exposed to the peanut and TMT odor swabs. We classified subtypes of *Pnoc*^BNST^ neurons based on individual response dynamics to peanut oil and TMT swabs during the first 10-seconds following the first entry into the near zone **(Figure 4A)**. We found that the proportion of *Pnoc*^BNST^ neurons that exhibited excited (31% vs. 30%), had no change (37% vs. 49%), or were inhibited (32% vs. 21%) during the initial exposure to the odor were similarly distributed across peanut and TMT odors **(Figure 4B)**. Interestingly we found that even though the proportion of excited neurons was similar with both peanut and TMT odors, when we averaged the activity of all the excited neurons for each group, we observed a sustained increase in activity dynamics of excited *Pnoc*^BNST^ neurons with the TMT odor **(Figure 4C and 4F)**.

**Figure 4.**
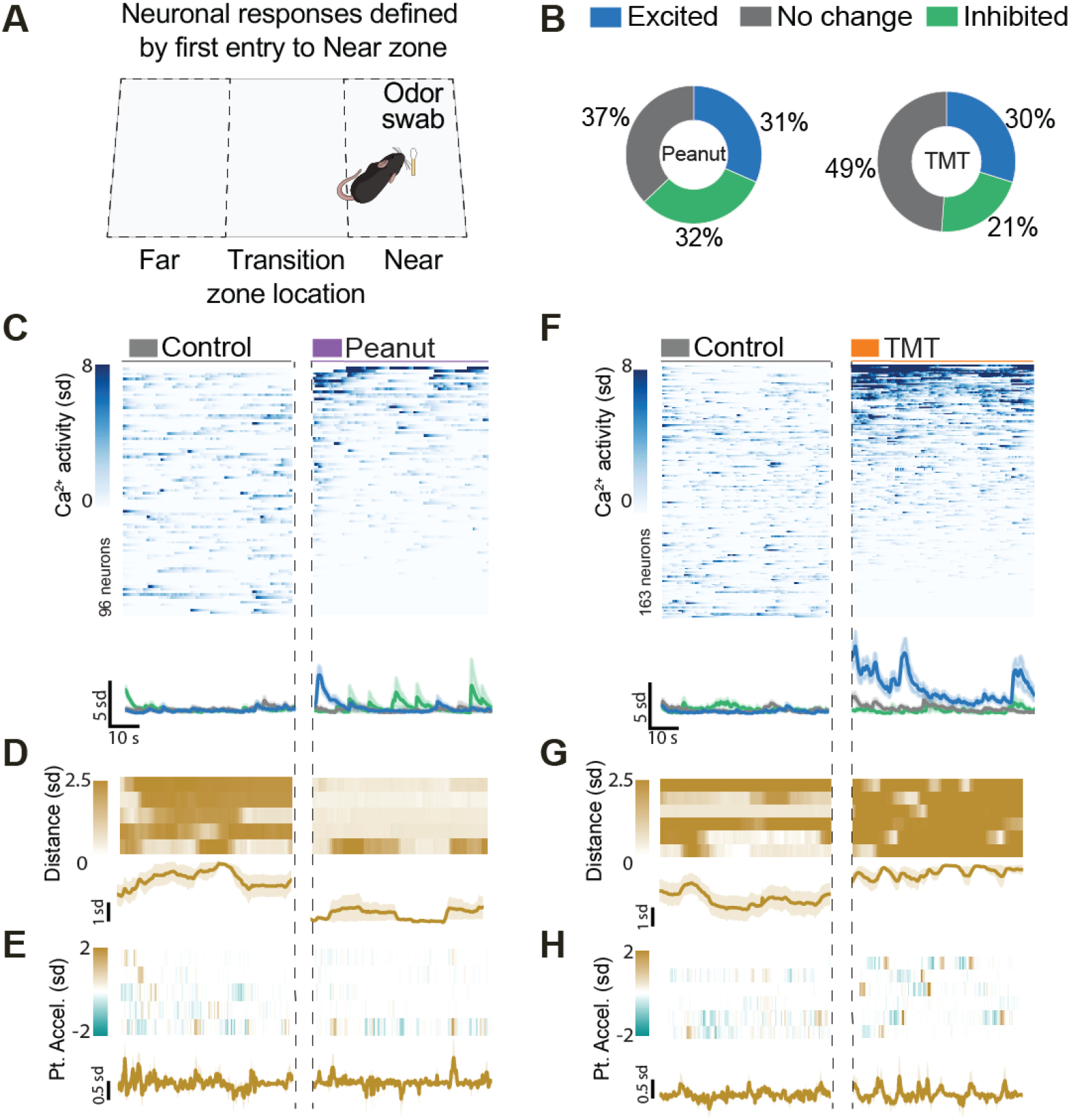
Identifying neurons based on individual response dynamics to TMT and Peanut odors. **(A)** Schematic of a freely moving mouse in its home cage to assess individual neuronal responses to water control, peanut oil swab, or TMT swab that are defined by first entry into the near zone. **(B)** Proportion of neurons exhibiting excitatory, no change, or inhibitory responses to peanut (n= 96 *Pnoc*^+^ neurons) and TMT odor (n=163 *Pnoc*^+^ neurons) upon first entry into the near zone. **(C, F)** Heatmaps of individual data (top) and grouped data (bottom) of *Pnoc*^BNST^ neuronal activity when mice are exposed to control, peanut oil, and TMT swab. **(D,E,G,F)** Distance and point acceleration of mouse to peanut (n=5 mice) and TMT odor (n=6 mice). Data is shown as mean ± SEM.

We next investigated differences in the distance between mice and the odor source when mice were exposed to either peanut or TMT odors. As expected, we found that the distance between mice and peanut odor was smaller compared to control odor **(Figure 4D)** and the distance between mice and TMT odor was larger compared to control **(Figure 4G)**. Lastly, we quantified point acceleration by measuring the changes in acceleration towards or away (ie. darting behavior) from the odor source. We found that mice showed more instances of darting away from the TMT odor compared to peanut odor **(Figure 4E and 4H)**.

### Excited *Pnoc*^BNST^ neurons correlate with darting away from TMT

Since we observed increased instances of darting behavior with mice that were exposed to TMT, we wanted to test if this behavioral action could be associated with the activity of individual *Pnoc*^BNST^ neurons. Therefore, we first analyzed if neuronal ensembles that exhibited either an excited, no change, or inhibited response to the initial encounter (10 s) with peanut or TMT odors correlated with distance to that particular odor. We found that neuronal ensembles that exhibited either an excited, no change, or inhibited response did not correlate with distance in mice exposed to the peanut odor **(Figure 5A)**. In contrast, individual *Pnoc*^BNST^ neurons that were excited to the initial exposure to TMT negatively correlated with distance (the closer ‘less distance’ the more activity, **Figure 5A)**. Next, we assessed if the activity dynamics of individual *Pnoc*^BNST^ neurons correlated with darting behavior, defined by increases in acceleration towards or away from the odor source (point acceleration). We found that neuronal ensembles that exhibited either an excited, no change, or inhibited response did not correlate with distance in mice exposed to the peanut odor **(Figure 5B)**. In contrast, individual *Pnoc*^BNST^ neurons that were excited to the initial exposure to TMT positively correlated with distance (the higher the point acceleration away from odor source the more activity, **Figure 5B)**. Together, these results indicate that individual response dynamics of *Pnoc*^BNST^ neurons signal both proximity towards and darting away from aversive odors.

**Figure 5.**
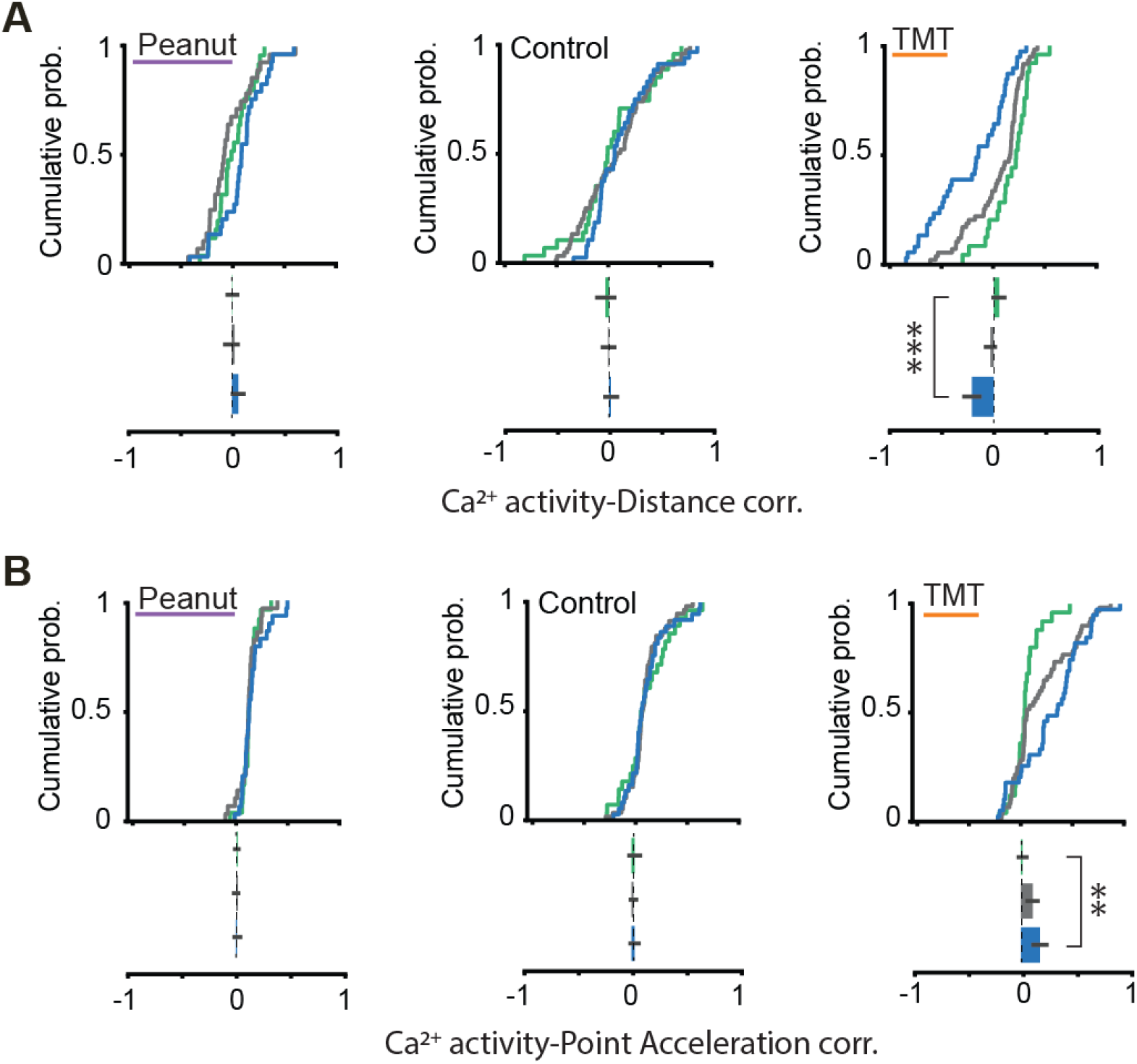
Excited *Pnoc*^BNST^ neurons correlate with proximity and darting away from TMT. **(A)** Correlation between Ca^2+^ activity dynamics of single *Pnoc*^BNST^ neurons and distance to odor source when mice were exposed to either water, peanut oil, or TMT odor swab (n= 96 *Pnoc*^+^ neurons from peanut odor group, n=163 *Pnoc*^+^ neurons from TMT odor group). **(B)** Correlation between Ca^2+^ activity dynamics of single *Pnoc*^BNST^ neurons and point acceleration towards or away from odor source when mice were exposed to either water, peanut oil, or TMT odor swab (n= 96 *Pnoc*^+^ neurons from peanut odor group, n=163 *Pnoc*^+^ neurons from TMT odor group). Data are shown as mean ± SEM. *\*\*p<0.01, ***p<0.001*.

## DISCUSSION

Dysregulated physiological arousal responses and disturbed motivated behavior are at the center of several neuropsychiatric disorders (Wilhelm and Roth, 2001; Maclean and Datta, 2007; Craske et al., 2009; Lang and McTeague, 2009; Schmidt et al., 2017; Urbano et al., 2017; Patriquin et al., 2019). However, these studies aimed to elucidate the neural correlates that underlie psychiatric disorders by investigating these two deleterious symptoms independently. Recently, we identified a subpopulation of neurons in the BNST (*Pnoc*^BNST^ neurons) that encode arousal responses to motivationally salient stimuli in head-fixed restrained mice (Rodriguez-Romaguera et al., 2020). The BNST is of high interest with regard to psychiatric illnesses due to its anatomical connectivity with brain regions that have an established role in motivation, arousal, reward, and fear (Lebow and Chen, 2016). Therefore, in the present study, we tested if arousal encoding *Pnoc*^BNST^ neurons play a significant role in encoding the motivated behavioral responses guided by stimuli of opposing valence. We find that the activity of a group of *Pnoc*^BNST^ neurons increases when mice are located near an aversive odor correlates with proximity to an aversive odor and darting away from the same odor. In addition, we also find that inhibiting the activity of *Pnoc*^BNST^ neurons did not reduce the motivation of mice to avoid an aversive odor. Lastly, we report that *Pnoc*^BNST^ neurons do not encode or signal behaviors to a rewarding food odor. Together, our findings reveal that the activity of *Pnoc*^BNST^ neurons is not necessary to maintain avoidance behavior, but signals the darting action that occurs when a mouse explores an aversive stimulus.

The results from our optogenetic experiment was initially surprising since we previously found that when we activate or inhibit the activity of *Pnoc*^BNST^ neurons in an elevated plus maze (EPM) assay, we can drive or attenuate anxiety-like behaviors, respectively (Rodriguez-Romaguera et al., 2020). In addition, these results contrast with another study that showed that inactivation of BNST reduced TMT-induced fear-related behavior (Fendt et al., 2003). The BNST is composed of several genetically identifiable cell-types (Giardino and Pomrenze, 2021; Ortiz-Juza et al., 2021), therefore, one explanation for why our inactivation of one cell-type in the BNST did not reduce TMT fear-related behavior could be that a different cell-type is responsible for TMT induced fear-related behavior. Another possible explanation for our finding could be that bulk inhibition of *Pnoc*^BNST^ neurons is not selective enough to modulate behavior. For instance, our single-cell analysis showed diversity in response dynamics within individual *Pnoc*^BNST^ neurons during exposure to both peanut and TMT odors. We previously found using single-cell sequencing that *Pnoc*^BNST^ neurons co-express other genetic markers such as somatostatin (*Som*), protein kinase C δ (*Pkcδ*), cholecystokinin (*Cck*) (Rodriguez-Romaguera et al., 2020). Each of these genetic markers anatomically cluster in distinct nuclei within the BNST (Ortiz-Juza et al., 2021), therefore, it is possible that the diverse response in *Pnoc*^BNST^ activity dynamics could be attributed to subtypes within the *Pnoc*^BNST^ neuronal population. Thus, bulk inhibition of the collective *Pnoc*^BNST^ neuronal population may not have been sufficient to modulate the specific neuronal ensemble responsible for driving the motivated behavior to aversive stimuli.

Furthermore, *Pnoc*^BNST^ neurons send dense projections into the medial preoptic area (mPOA) and into the medial amygdala (MeA) (Rodriguez-Romaguera et al., 2020). How *Pnoc*^BNST^ neurons influence the downstream neural activity within these two regions is unknown. The MeA is a region studied for fear-related behaviors (Fendt et al., 2003; Li et al., 2004; Müller and Fendt, 2006; Takahashi et al., 2007; Arakawa et al., 2010; McCue et al., 2014; Miller et al., 2019). The inhibition of MeA via local infusion of the GABA_A_ agonist muscimol, reduced TMT-induced fear behavior (Müller and Fendt, 2006). *Pnoc*^BNST^ neurons are primarily GABAergic, therefore, it is possible that optically targeting the specific *Pnoc*^BNST^ neurons that project into the MeA would modulate motivated behavior to TMT. This remains to be tested, however, the result from this study might suggest a role of *Pnoc*^BNST^ neuron activity in defensive behaviors via downstream communication with MeA.

Here, we used advanced tools that allowed us to track the activity of *Pnoc*^BNST^ neurons with single-cell resolution in unrestrained mice that freely behave. We discovered that a subset of *Pnoc*^BNST^ neurons signal darting away from aversive stimuli. However, to test if this ensemble of *Pnoc*^BNST^ neurons is necessary for darting behavior, we need to employ advanced holographic optogenetic techniques with the ability to precisely target custom neural ensembles, with high spatial (Pégard et al., 2017; Xue et al., 2022) and temporal resolution (Eybposh et al., 2020; Curtis et al., 2021) to modulate the activity dynamics of these functionally defined ensembles of *Pnoc*^BNST^ neurons that signal darting behavior. Lastly, to further investigate *Pnoc*^BNST^ neurons in arousal and motivation in freely moving animals, future studies will need to employ strategies that can track rapid changes in physiological arousal simultaneously with *in vivo* calcium imaging. Thus far, techniques to track pupillometry in freely moving mice are limited and prevent the ability to employ freely moving calcium imaging strategies. As the field of pupillometry expands, we will be able to refine our understanding of arousal during ongoing motivated behaviors and how both of them contribute to the development of neuropsychiatric disorders.

## ACKNOWLEDGEMENTS

This work was supported by grants from the National Institute of Mental Health (F30-MH115693, R.L.U.), the National Science Foundation (DGE-1650116, M.M.O.-J.), the UNC Royster Fellowship (V.R.C.), the Helen Lyng White Fellowship (G.V.-H.), Nationa Institute of General Medical Sciences: UNC’s PREP in the Biomedical Sciences (R25GM089569, R.A.G.-R.), the National Institute of Drug Abuse (R37-DA032750 and R01-DA038168, G.D.S.), the Brain and Behavior Research Foundation (G.D.S.), the National Institute on Deafness and Other Communications Disorders (R01-DC017516, H.K.K.), the Burroughs Wellcome Fund (N.C.P), the Arnold and Mabel Beckman Foundation (N.C.P), the National Institute of Mental Health (F32-MF113327, J.R.-R.), the Foundation of Hope (G.D.S., H.K.K., J.R.-R.), a Junior Faculty Development Award from the UNC Provost's Office, sponsored by IBM and R.J. Reynolds (J.R.-R.), and the UNC Department of Psychiatry (J.R.-R.).

## MATERIAL AND METHODS

Detailed methods are provided in the online version of this paper and include the following:

- KEY RESOURCES TABLE
- STATISTICAL ANALYSIS
- CONTACT FOR REAGENTS AND RESOURCE SHARING
- EXPERIMENTAL MODEL AND SUBJECT DETAILS

- Animals
- METHOD DETAILS

- Viral Constructs
- Surgical Procedure and Histology
- Odor Preference in Home Cage
- Calcium Imaging in Freely moving Mice
- Optogenetic Manipulations
- QUANTIFICATION AND STATISTICAL ANALYSIS

- Behavioral Optogenetics Data Analysis
- Calcium Imaging Analysis

DATA AND SOFTWARE AVAILABILITY

### KEY RESOURCES TABLE (TABLE S1)

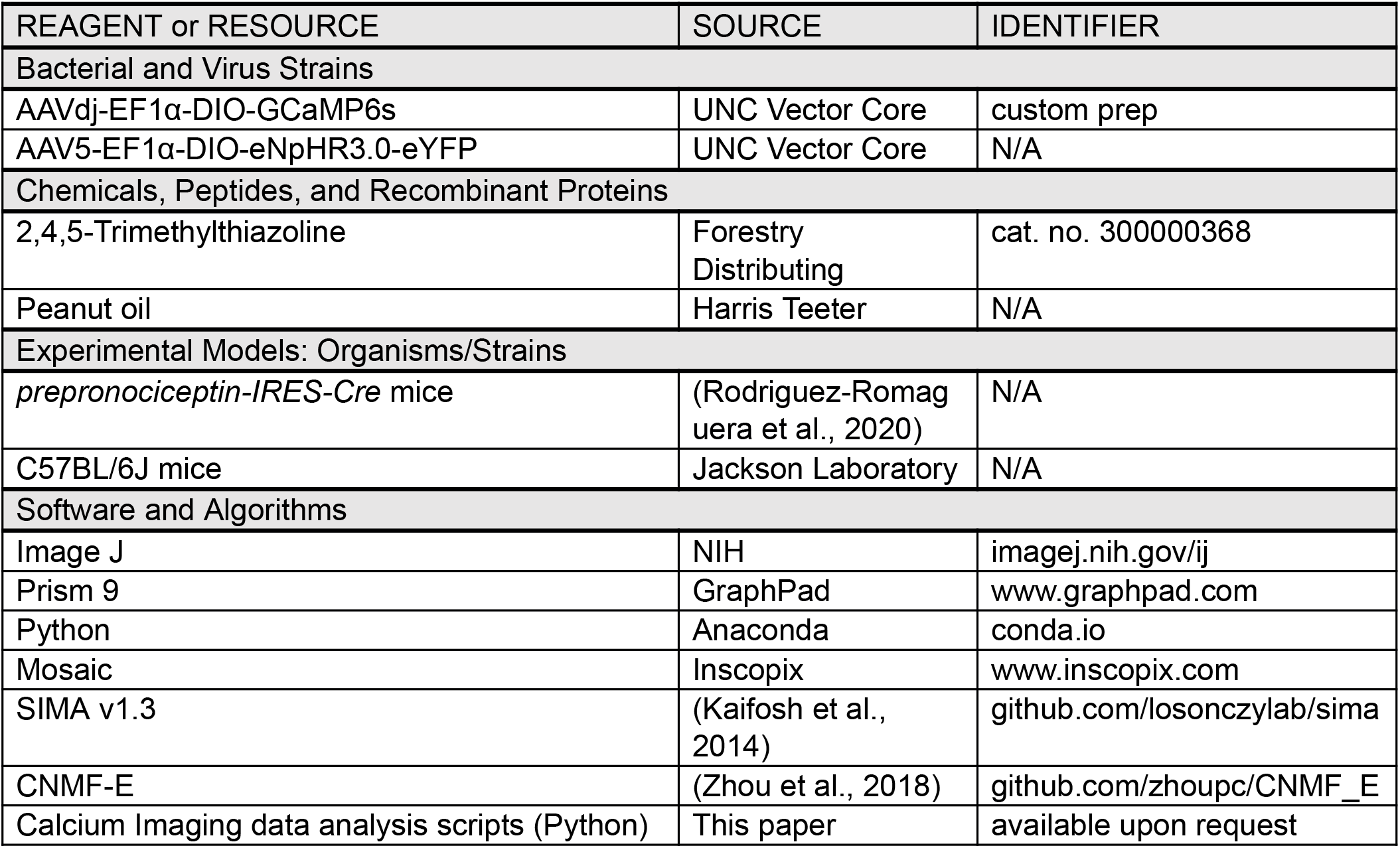

### STATISTICAL ANALYSIS

| Figure | # of Samples | Test | t, F, U Value | p Value |
| --- | --- | --- | --- | --- |
| **Figure 1** |  |  |  |  |
| C; Percent time in Peanut odor zones compared to water control odor. | n = 5 *Pnoc-Cre* mice | Paired t-test, no correction | <u>Near zone:</u><br>$t = -3.03$<br><u>Transition zone:</u><br>$t = 1.47$<br><u>Far zone:</u><br>$t = 2.99$ | $p=0.039$<br>$p=0.215$<br>$p=0.040$ |
| D; Percent time in TMT odor zones compared to water control odor. | n = 6 *Pnoc-Cre* mice | Paired t-test, no correction | <u>Near zone:</u><br>$t = 2.68$<br><u>Transition zone:</u><br>$t = -0.08$<br><u>Far zone:</u><br>$t = -2.41$ | $p=0.044$<br>$p=0.941$<br>$p=0.061$ |
| **Figure 2** |  |  |  |  |
| F; Comparison of average $Ca^{2+}$ activity of control water swab and peanut swab by odor zones. | n = 96 *Pnoc*<sup>+</sup> neurons from 4 *Pnoc-Cre* mice | Two-way ANOVA test<br>Bonferroni's multiple comparisons | <u>Zone:</u><br>$F_{(1,8)}=0.1874$<br><u>Odor:</u><br>$F_{(1,8)}=0.02448$<br><u>Interaction:</u><br>$F_{(1,8)}=0.4894$<br><u>Near zone:</u><br>$t = 0.7588$<br><u>Transition zone:</u><br>$t = 0.3288$<br><u>Far zone:</u> n/a | $p=0.6766$<br>$p=0.8795$<br>$p=0.5040$<br>$p=0.9395$<br>$p=>0.9999$ |
| G; Comparison of average $Ca^{2+}$ activity of control water swab and TMT swab by odor zones. | n = 163 *Pnoc*<sup>+</sup> neurons from 6 *Pnoc-Cre* mice | Two-way ANOVA test<br>Bonferroni's multiple comparisons | <u>Zone:</u><br>$F_{(2,23)}=0.8205$<br><u>Odor:</u><br>$F_{(1,23)}=16.65$<br><u>Interaction:</u><br>$F_{(2,23)}=0.5628$<br><u>Near zone:</u><br>$t = 3.018$<br><u>Transition zone:</u><br>$t = 2.244$<br><u>Far zone:</u><br>$t = 1.768$ | $p=0.4527$<br>$p=0.0005$<br>$p=0.5772$<br>$p=0.0184$<br>$p=0.1042$<br>$p=0.2711$ |
| H; Comparison of average $Ca^{2+}$ activity of Peanut and TMT | n = 96 *Pnoc*<sup>+</sup> neurons from 4 *Pnoc-Cre* mice<br>n = 163 *Pnoc*<sup>+</sup> neurons from 6 *Pnoc-Cre* mice | Bonferroni's multiple comparisons | <u>Water - PNT</u><br>$t= 0.6411$<br><u>Water - TMT</u><br>$t=4.538$ | $p>0.9999$<br>$p=0.0038$ |

| **Figure 3** |  |  |  |  |
| --- | --- | --- | --- | --- |
| D; Time near TMT odor zone | n = 7 NpHR-Cre<sup>-</sup> mice<br>n = 5 NpHR-Cre<sup>+</sup> mice | Two-way ANOVA test<br>Bonferroni | Group:<br>$F(1,10)=0.5271$<br>Laser:<br>$F(1,10)=13.08$<br>Interaction:<br>$F(1,10)=0.1066$<br>Laser Off<br>Laser On | $p=0.4845$<br>$p=0.0047$<br>$p=0.7486$<br>$p=0.8698$<br>$p=0.9999$ |
| **Figure 4** |  |  |  |  |
| B; Proportion of neurons excited, inhibited, or showed no change in response upon first entry of the near odor zone. | n = 96 *Pnoc*<sup>+</sup> neurons for PNT odor<br>n= 163 *Pnoc*<sup>+</sup> neurons for TMT odor | Chi-squared | $\chi^2(2) = 3.974$ | $p=0.14$ |
| **Figure 5** |  |  |  |  |
| A; Correlation between Ca<sup>2+</sup> activity responses and distance when mice were exposed to peanut odor. | n = 96 *Pnoc*<sup>+</sup> neurons for PNT odor | ANOVA; Bonferroni corrected | <u>Inhibited vs No Change:</u><br>$t= 0.7019$<br><u>Excited vs No Change:</u><br>$t= 2.229$<br><u>Excited vs. Inhibited:</u><br>$t= 1.456$ | $p>0.9999$<br>$p=0.0860$<br>$p=0.4479$ |
| Correlation between Ca<sup>2+</sup> activity responses and distance when mice were exposed to TMT odor. | n= 163 *Pnoc*<sup>+</sup> neurons for TMT odor | | <u>Inhibited vs No Change:</u><br>$t= 1.701$<br><u>Excited vs No Change:</u><br>$t= 4.643$<br><u>Excited vs. Inhibited:</u><br>$t= 5.308$ | $p=0.2746$<br>$p=<0.0001$<br>$p=<0.0001$ |
| B; Correlation between $\text{Ca}^{2+}$ activity responses and point acceleration when mice are exposed to peanut odor. | n = 96 *Pnoc*<sup>+</sup> neurons for PNT odor | ANOVA; Bonferroni corrected | <u>*Inhibited vs No Change:*</u><br>*t*= 0.3233<br><u>*Excited vs No Change:*</u><br>*t*= 1.449<br><u>*Excited vs. Inhibited:*</u><br>*t*= 1.065 | *p*>0.9999<br>*p*=0.4540<br>*p*=0.8700 |
| Correlation between $\text{Ca}^{2+}$ activity responses and point acceleration when mice are exposed to TMT odor. | n= 163 *Pnoc*<sup>+</sup> neurons for TMT odor | | <u>*Inhibited vs No Change:*</u><br>*t*= 2.185<br><u>*Excited vs No Change:*</u><br>*t*= 1.789<br><u>*Excited vs. Inhibited:*</u><br>*t*= 3.466 | *p*=0.0925<br>*p*=0.2282<br>*p*=0.0022 |

### CONTACT FOR REAGENTS AND RESOURCE SHARING

Information and requests for reagents may be directed and will be fulfilled by the corresponding author Jose Rodriguez-Romaguera.

### EXPERIMENTAL MODEL AND SUBJECT DETAILS

#### Animals

Adult (25-30g) male *prepronociceptin*-IRES-*Cre* (*Pnoc*-IRES-Cre) or wild type mice (C57BL6/J) were independently housed and maintained on a reverse 12-hr light-dark cycle (lights off at 08:00 AM) with *ad libitum* access to food and water. Behavior was tested during the dark cycle. All procedures were conducted in accordance with the Guide for the Care and Use of Laboratory Animals, as adopted by the National Institute of Health, and with the approval of the Institutional Animal Care and Use Committee from the University of North Carolina at Chapel Hill.

### METHOD DETAILS

#### Viral Constructs

Both Cre-inducible viral constructs [AAVdj-EF1α-DIO-GCaMP6s (3.1 × 10^12^ infectious units/mL) and AAV5-EF1α-DIO-eNpHR3.0-eYFP (8.0 × 10^12^ infectious units/mL)] were packaged by the UNC Vector Core.

#### Surgical Procedure and Histology

Mice were anesthetized with isoflurane (0.8-1.5%) vaporized in pure oxygen (1 l/min^−1^) and placed in a stereotaxic frame (David Kopf Instruments, Tujunga, CA). Ophthalmic ointment (Akorn, Lake Forest, IL), a topical anesthetic (2% lidocaine; Akorn, Lake Forest, IL), and triple antibiotic ointment (NWI, Athens, TN) were applied during surgeries. Mice received a subcutaneous injection of saline (0.9% NaCL in water) to offset fluid loss during surgery and an injection of Meloxicam (0.5 mg/kg) for pain relief. A 33-gauge injection needle connected to a 2 uL syringe (Hamilton Company, Reno, NV) delivered viruses into the anterior dorsal portion of the bed nucleus of the stria terminalis (adBNST; 500 nl per side; relative to bregma: +0.14 AP, +/-0.95 ML, DV −4.20 DV). For calcium imaging studies, unilateral virus injections were made into adBNST followed by implantation of a gradient refractive index lens [GRIN lens; Inscopix, Palo Alto, CA (0.6 mm in diameter, 7.3 mm in length)] placed 0.3 mm dorsal to adBNST target site. To image BNST neurons in freely moving mice, a magnetic baseplate was cemented around the GRIN lens (Ghosh et al., 2011; Resendez et al., 2016) to secure the attachment of a miniaturized head-mounted microscope (nVista 2.0, Inscopix, Palo Alto, CA). For optogenetic experiments, bilateral virus injections were made into adBNST, and an optical fiber was implanted with a 10° angle approximately 0.5 mm above adBNST. Following surgery, mice were given acetaminophen in their drinking water for two days for pain relief.

After behavioral experiments, all cohorts were deeply anesthetized with isoflurane and transcardially perfused with 1X PBS followed by ice-cold 4% paraformaldehyde solution (pH 7.4, Thermo Fisher Scientific, Waltham, MA). Brain tissue was extracted and placed overnight in 4% paraformaldehyde solution at 4°C then moved into a 30% sucrose solution for two days. Coronal brain sections (40 μm) were collected with a cryostat, counterstained with DAPI and coverslipped to verify viral expression and optic fiber or GRIN lens placement.

#### Odor Preference in Home Cage

Mice habituated to a square block holder in their home cage for two days prior to testing. On the day of testing, a cotton swab was placed in the square block holder located in an upright position 4 inches from the home cage floor on one of the sides (sides were alternated across mice). Behavior was recorded for a 5-min period after placing 2.5 μl of distilled H_2_O on the cotton swab, followed by placing either 2.5 μl of trimethylthiazoline (TMT) or 2.5 μl of peanut oil (Harris Teeter, Carrboro, NC) on the cotton swab. Distance to odor (cm, max: 25 cm), time spent freezing (s), and velocity (cm/s) were calculated using automated tracking software (Ethovision XT 11, Noldus, Leesburg, VA). A low dose of TMT was used to minimize freezing responses and maintain ambulation. Additionally, mice were not exposed to both TMT and peanut oil to avoid learned association between the cotton swab and a given salient odor.

#### Calcium Imaging in Freely Behaving Mice

A miniaturized head-mounted microscope (nVista 2.0, Inscopix, Palo Alto, CA) was used to visualize activity dynamics of *Pnoc*^+^ neurons in BNST *in vivo* and in freely moving animals. To achieve such, a virus encoding the Cre-dependent calcium indicator GCaMP6s (AAVdj-EF1α-DIO-GCaMP6s; 3.1 × 10^12^ infectious units/mL) was injected into the BNST of *Pnoc-Cre-IRES* mice (see Surgical Procedure and histology section). After a minimum of 8 weeks to allow sufficient time for virus transport and infection, mice were habituated to a tethered dummy microscope during 1-hour sessions across 3 consecutive days. Each day the microscope was attached to the baseplate mounted on the cranium while mice were under light anesthesia (1.5 % isoflurane). During testing, mice were given 45 minutes to recover from anesthesia before being exposed to the odor preference assay during which GCaMP6s-expressing neurons were visualized. Using nVista HD Acquisition Software (Inscopix, Palo Alto, CA), images were scanned at 15 frames per second with the LED transmitting 0.1 to 0.2 mW of light on average. Videos were motion-corrected using Mosaic software (Inscopix, Palo Alto, CA) and then downsampled to 5 frames per second. Calcium transients and deconvolved events were extracted from individual ROI’s using a version of CNMF that was adapted for one-photon miniature microscope video data (CNMF-E) (Zhou et al., 2018) and data was analyzed using custom data analysis pipelines written in Python (see Quantification and Statistical Analysis section).

#### Optogenetics

Photoinhibition manipulation experiments were performed as previously described (Rodriguez-Romaguera et al., 2020). A virus encoding the Cre-inducible halorhodopsin (AAV5-ef1α -eNpHR3.0-eYFP; 8.0 × 10^12^ infectious units per ml) was injected into BNST of either *PNOC-Cre-IRES* mice or their wild type littermates as controls. Behavior experiments proceeded after ≥ 4 weeks to allow for adequate virus transduction. Prior to behavioral testing, mice were habituated to an optic fiber tether in their home cage for 3 days. On testing day, mice were placed into their home cage with a TMT odor source located on one side (this in a similar fashion described in Odor preference in the home cage). The laser (532 nm; 8-10 mW) was constantly applied during a 3 min period followed by a 3 min laser off period that repeated for a total of 18 minutes.

### QUANTIFICATION AND STATISTICAL ANALYSIS

#### Behavioral and Optogenetics Data Analysis

The data obtained from the odor preference and optogenetic experiments were analyzed using Prism 9 (GraphPad Software Inc., La Jolla, CA). Mean values are accompanied by SEM values. Comparisons were tested using paired or unpaired t-tests. Two-way ANOVA tests followed by either Tukey’s post-hoc tests or Bonferroni post-hoc comparisons were applied for comparisons with more than two groups, n.s. p > 0.05, ^#^p <0.1 (to signify a trend), *p < 0.05, **p < 0.01, ***p < 0.001.

#### Calcium Imaging Analysis

Calcium imaging recordings were first motion-corrected using a planar hidden Markov model (Kaifosh et al., 2014). Neurons were identified, and their calcium signals were extracted using an implementation of constrained nonnegative matrix factorization adapted for micro-endoscopic data (CNMF-E) (Zhou et al., 2018), which accounts for the unusually large background fluctuations and demixes the spatially overlapping signals. This extracted signal was scaled to account for variations in fluorescence intensities among cells by the standard deviation of a neuron’s fluorescence throughout an experiment. For home cage, odor-presentation experiments, neurons were defined as excited to TMT or Peanut Oil by comparing fluorescence values for frames between that odor and water (the control) using a Mann-Whitney U test with Bonferroni correction. Processing after signal extraction was performed using algorithms written in Python.

### DATA AND SOFTWARE AVAILABILITY

Python was used to analyze all the calcium imaging datasets (written by R.L.U.). The codes used for this analysis are available upon request to the corresponding author.

